# Identification of Reptarenaviruses, Hartmaniviruses and a Novel Chuvirus in Captive Brazilian Native Boa Constrictors with Boid Inclusion Body Disease

**DOI:** 10.1101/2020.01.02.893420

**Authors:** Fernando Froner Argenta, Jussi Hepojoki, Teemu Smura, Leonora Szirovicza, Márcia Elisa Hammerschmitt, David Driemeier, Anja Kipar, Udo Hetzel

**Author notes:** The authors contributed equally to this manuscript, the author order was determined alphabetically (Fernando Froner Argenta, Jussi Hepojoki). **Corresponding author:** Present address: University of Helsinki, Medicum, Department of Virology, Haartmaninkatu 3, FI-00290 Helsinki, Finland, or.

## Abstract

Boid Inclusion Body Disease (BIBD) is a transmissible viral disease of captive snakes that causes severe losses in snake collections worldwide. It is caused by reptarenavirus infection, which can persist over several years without overt signs, but is generally associated with the eventual death of the affected snakes. Thus far, reports have confirmed existence of reptarenaviruses in captive snakes in North America, Europe, and Australia, but there is no evidence that it also occurs in wild snakes. BIBD affects both boas and pythons, the habitats of which do not naturally overlap. Herein, we studied Brazilian captive snakes with BIBD using a metatranscriptomic approach, and report the identification of novel reptarenaviruses, hartmaniviruses, and a new species in the family *Chuviridae*. The reptarenavirus L segments identified represent six novel species, while we only found a single novel reptarenavirus S segment. Until now, hartmaniviruses had been identified only in European captive boas with BIBD, and the present results increase the number of known hartmanivirus species from four to six. The newly identified chuvirus showed 38.4%, 40.9%, and 48.1% amino acid identity to the nucleoprotein, glycoprotein, and RNA-dependent RNA polymerase of its closest relative, Guangdong red-banded snake chuvirus-like virus. Although we cannot rule out the possibility that the found viruses originated from imported snakes, the results suggest that the viruses would circulate in indigenous snake populations.

**IMPORTANCE:** Boid Inclusion Body Disease (BIBD) caused by reptarenavirus infection affects captive snake populations worldwide, but the reservoir hosts of reptarenaviruses remain unknown. Herein, we report the identification of novel reptarenavirus and hartmanivirus species, and a chuvirus in captive Brazilian boas with BIBD. Three of the four snakes studied showed co-infection with all three viruses, and one of the snakes harbored three novel reptarenavirus L and one novel S segment. The samples originated from collections with Brazilian indigenous snakes only, which could indicate that these viruses circulate in wild snakes. The findings could further indicate that boid snakes are the natural reservoir of reptarena- and hartmaniviruses commonly found in captive snakes. The snakes infected with the novel chuvirus all suffered from BIBD; it is therefore not possible to comment on its potential pathogenicity and contribution to the observed changes in the present case material.

## INTRODUCTION

The global decline in biodiversity is a topic of concern also for members of the class Reptilia. The worldwide transportation of wild caught, farm- and captive-bred reptiles facilitates also the transmission of pathogens. Thus, further information on reptilian pathogens is required to enable efficient screening of transported animals in order to secure e.g. zoological collections and to avoid spread of infectious agents into private and commercial breeding collections. Boid Inclusion Body Disease (BIBD), known to affect captive constrictor snakes, was recognized in the 1970s (1, 2), and arenaviruses were identified as the causative agent(s) in the early 2010s (3–10). BIBD affects nonvenomous constrictor snakes inhabiting biotopes in the neotropics and tropics. The natural habitats of boas include Central and South America, and Madagascar, while pythons are inherent in Africa, Asia and Australia. Although the habitats of boas and pythons do naturally not overlap geographically, snake species from several continents are housed together or in close proximity in zoological and private collections all around the world. As the name implies, BIBD manifests by the formation of eosinophilic and electron-dense inclusion bodies (IBs) within almost all cell types (2, 3, 5, 11). In fact, the ante mortem BIBD diagnosis relies on the detection of IBs in cytological specimens, e.g. blood smears (12, 13), or liver biopsies (1, 14). The identification of reptarenaviruses as the causative agent for BIBD has enabled RT-PCR based diagnostic procedures and screening of collections (12, 13, 15). Due to reasons unknown, BIBD is diagnosed more often in boas than in pythons (1, 10, 14). The disease can manifest itself with central nervous system (CNS) signs, which include opisthotonus (“star-gazing”), head tremors, disorientation, regurgitation and “corkscrewing” (1, 2). However, during the past decades, boas with BIBD and clinical CNS signs have become rare and even snakes with extensive IB formation often appear clinically healthy (10, 12, 14), which could be an indication of adaptation towards lower virulence. Instead, snakes with BIBD seem to emaciate progressively and become terminally ill due to secondary, usually bacterial infections, presumably due to BIBD-associated immunosuppression (13).

In 2015, the BIBD associated arenaviruses were grouped to form the genus *Reptarenavirus* in the family *Arenaviridae*, and the formerly known arenaviruses of rodents and bats formed the genus *Mammarenavirus* (16). The mamm- and reptarenavirus genome is a bisegmented negative-sense RNA with ambisense coding strategy (17). The S segment encodes the glycoprotein precursor (GPC) and nucleoprotein (NP), and the L segment encodes the zinc finger matrix protein (ZP) and the RNA-dependent RNA polymerase (RdRp) (17). Coincidentally, we identified Haartman Institute Snake virus-1 (HISV-1) in a snake with BIBD (9), and later demonstrated that the genome of HISV-1 is similar to that of mamm- and reptarenaviruses, except that it lacks the ZP gene (18). The identification of HISV-1 led to the formation of a third arenavirus genus, *Hartmanivirus* (19). The most recent addition to the family *Arenaviridae* is the genus *Antennavirus*, the representatives of which carry three instead of two genome segments (20). Others and we have demonstrated that snakes with BIBD often show co-infection with several reptarenavirus species (8, 9). We also identified further hartmaniviruses and showed that hartmaniviruses can co-infect snakes with BIBD (18). However, so far it is not clear whether hartmaniviruses contribute to BIBD pathogenesis.

The origin of reptarenaviruses and hartmaniviruses is still unknown, as reports have only described BIBD diagnosed in captive snakes. However, in order to gather information whether boid snakes themselves can be the viral reservoirs, it is of particular interest to see whether BIBD occurs within boid snake populations in the natural habitats. *Boa constrictors* are indigenous in Brazil, and the knowledge on reptarenavirus occurrence is limited to a single case report of a suspected BIBD case in *Corallus annulatus* kept in a zoological garden (21). In 2017, we diagnosed the first cases of BIBD in captive Brazilian *Boa constrictor* and undertook the present study to investigate the nature and phylogeny of the involved causative viruses.

## RESULTS

### Case descriptions

#### Clinical histories

Animals # 1 and #4 died after unsuccessful therapeutic attempts (antibiotic and fluid therapy, catheter feeding) and chronic inflammatory processes in oral cavity and sinuses and a period of apathy, animal #3 died after a prolonged period of apathy and neurological signs, and animal #2 was found dead without prior clinical signs (Table 1).

**Table 1:** Animals and pathological findings. All animals were *Boa constrictor constrictor* snakes held in captivity in Porto Alegre, Brazil.

| Case No | Age | Sex | Origin | Captivity | Clinical history | Diagnoses |
| --- | --- | --- | --- | --- | --- | --- |
| [1] | 12 y | F | Novo Arão<br>(Amazonas,<br>Brazil) | Private<br>owner | Regurgitation and ulcerative<br>lesion in oral cavity for 3 mo.<br>Antibiotic therapy, fluid<br>therapy, catheter feeding for 2<br>mo; no improvement, then<br>apathy and death. | Chronic ulcerative stomatitis and<br>osteomyelitis (maxilla); chronic ulcerative<br>dermatitis and myositis; BIBD |
| [2] | 10 y | F | Novo Arão<br>(Amazonas,<br>Brazil) | Private<br>owner | Found dead without prior<br>clinical signs. | Fibrinonecrotizing cloacitis, embolic-metastatic nephritis; hepatic lipidosi-<br>s; BIBD |
| [3] | 12 y | M | Novo Arão<br>(Amazonas,<br>Brazil) | Private<br>owner | Apathy and neurological signs<br>(eg. disorientation, star-gazing) for 5 mo; euthanasia | Focally ulcerated granulomatous<br>stomatitis and osteomyelitis (maxilla);<br>BIBD |
| [4] | 13 y | M | Unknown | Botanic Zoo<br>Foundation<br>of Rio<br>Grande | Oral mucosal bleeding, nasal<br>discharge, lethargy, anorexia<br>for appr. 5 mo; hospitalization<br>and antibiotic therapy, fluid<br>therapy, catheter feeding; no<br>improvement; after 45 d<br>apathy and death. | Emaciation; chronic suppurative<br>sinusitis; BIBD |

#### Post mortem findings

At necropsy, animals #1-3 exhibited good body condition, whereas animal #4 was emaciated. All four snakes exhibited overt inflammatory processes: a chronic ulcerative stomatitis and osteomyelitis of the maxilla (animals #1 and #3) (Fig. 1), a multifocal ulcerative deep dermatitis and myositis extending to the vertebral bones (animal #1), a fibrino-necrotizing cloacitis (animal #2), and a chronic suppurative sinusitis (animal #4). Histological examination confirmed the findings. In animal #1, the stomatitis was predominantly heterophilic (i.e. suppurative); bacteriology and mycology isolated *Enterobacter gergoviae, Providencia ssp., Proteus ssp*., and *Candida albicans*, respectively, from the lesions. The inflammatory infiltrate of the cloacitis in animal #2 was heterophil-dominated, with abundant aggregates of coccoid bacilli within the superficial layer of fibrin and necrotic debris. This animal exhibited multifocal areas of necrosis with embedded aggregates of coccoid bacilli in the kidneys, consistent with embolic-metastatic nephritis as a consequence of bacteremia. A bacteriological examination was not performed. In animal #3, the stomatitis was granulomatous, with multinucleated giant cells; only non-specific flora *(Klebsiella ssp.)* was isolated from the lesion. In the sinusitis of animal #4 heterophils were the predominant inflammatory cells; the bacteriological examination of a sample from this lesion yielded exuberant mixed growth *(Enterobacter gergoviae* and *Pseudomonas* ssp.). In addition, this animal exhibited a mild multifocal heterophil-dominated pneumonia.

**Figure 1.**
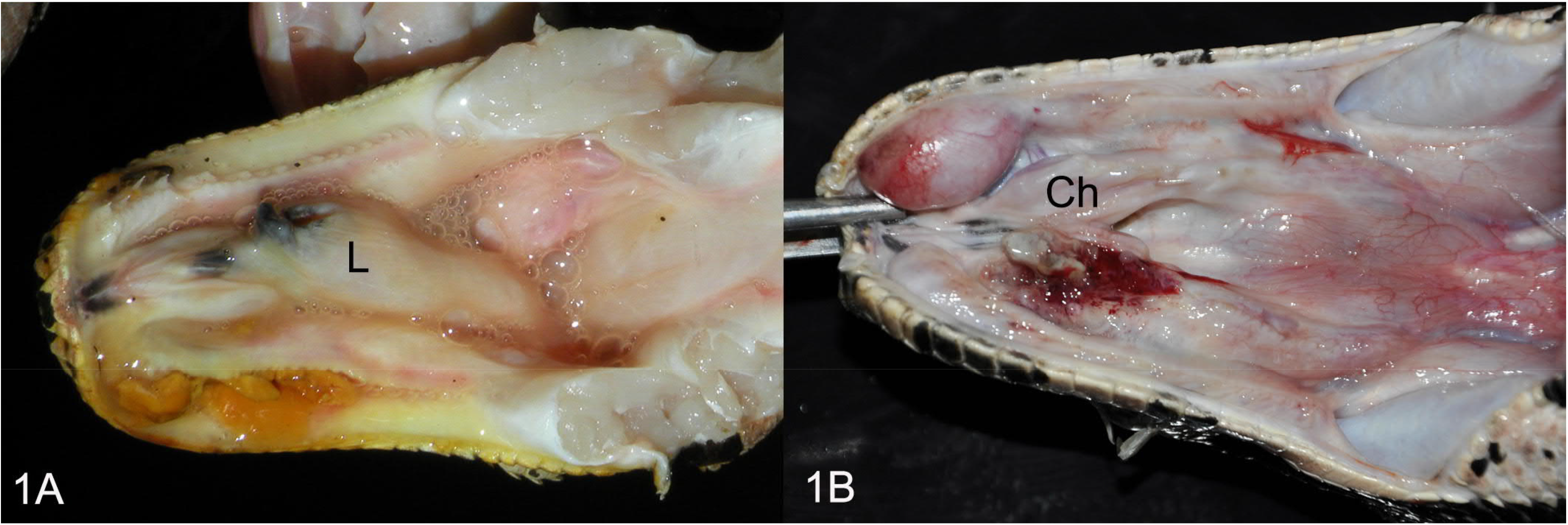
Oral cavity of snakes with confirmed BIBD. **A**. Animal #1. Chronic ulcerative stomatitis. L: larynx. **B**. Animal #3. Focally ulcerated chronic stomatitis. Ch: choana.

In all snakes, histology served to confirm BIBD. The characteristic eosinophilic intracytoplasmic IBs were found in parenchymal cells in a range of organs (brain (Figs. 2A) and spinal cord, liver (Fig. 3A), pancreas, lungs, kidneys) in all animals. The IBs varied in size distribution, indicating a chronic stage of the disease (Figs. 2A and 3A).

**Figure 2.**
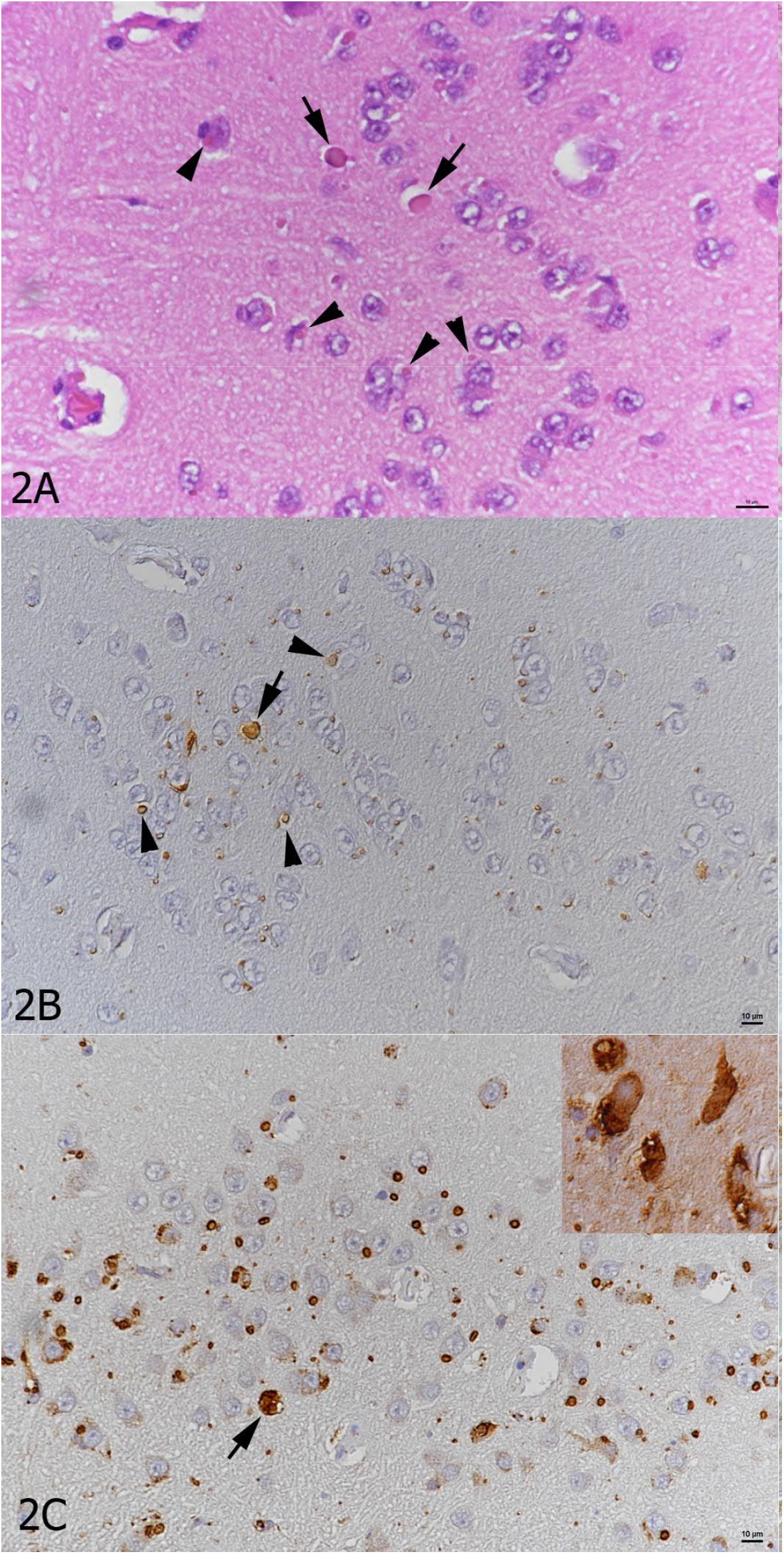
Histological and immunohistological findings in the brain of animal #3. **A**. Neurons exhibit the typical cytoplasmic eosinophilic IBs (arrowheads) which vary in size and can reach the size of and obscuring the nucleus (arrows). HE stain. **B**. Staining with the anti-pan-RAV antibody highlights the IBs depicted in the HE stain. Immunohistology, hemalaun counterstain. **C**, Staining with the anti-pan-hartmani antibody highlights the IBs, but also shows the presence of NP within the entire cytoplasm of infected cells (arrow and inset). Immunohistology, hemalaun counterstain.

**Figure 3.**
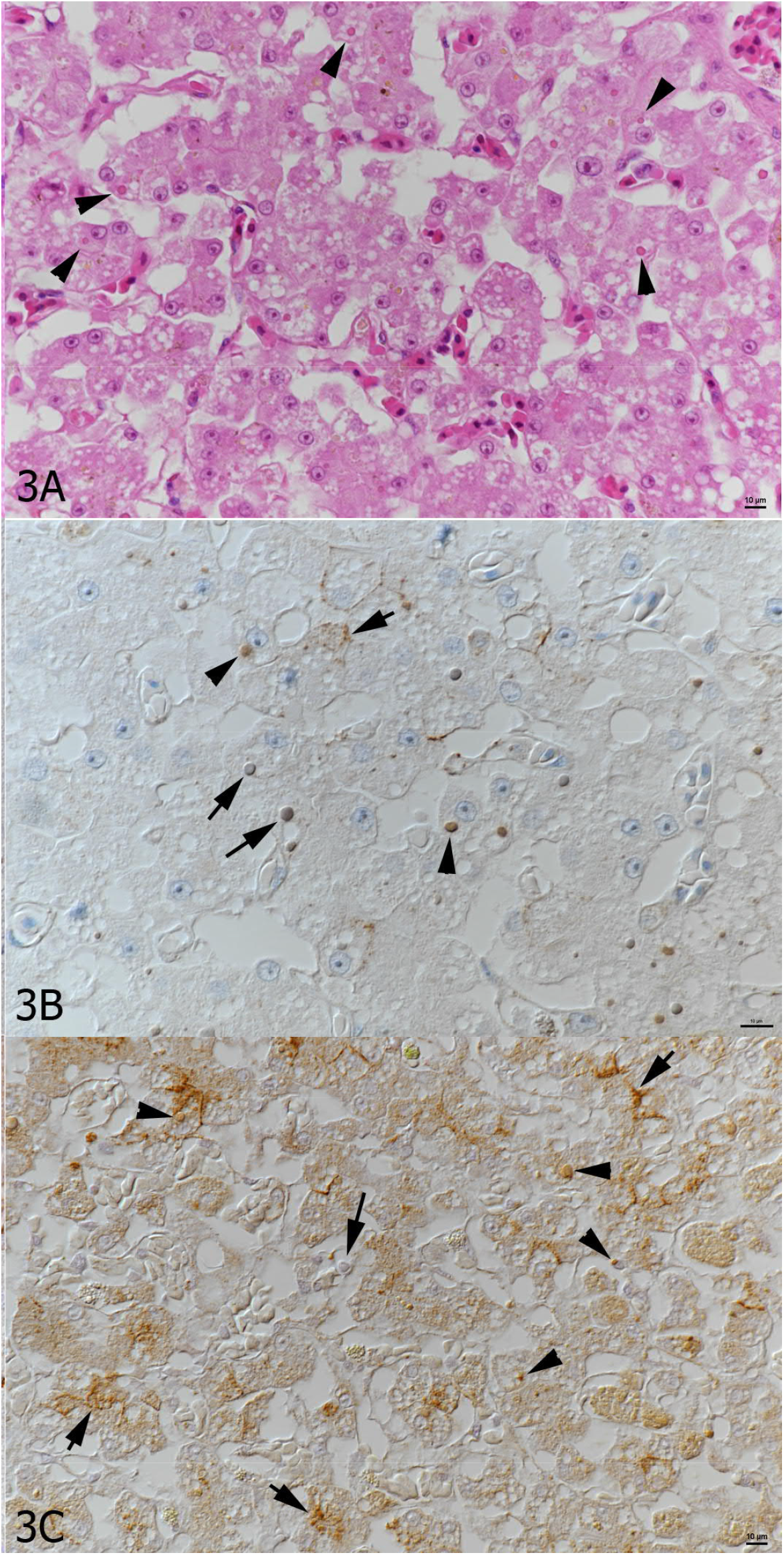
Histological and immunohistological findings in the liver of animal #4. **A**. Numerous hepatocytes exhibit a cytoplasmic eosinophilic IB of variable size (arrowheads). HE stain. **B**. Staining with the anti-UHV NP antibody highlights individual IBs (arrowheads) and shows that some cells contain several small IBs (short arrow). Some larger IBs appear negative (large arrows). Immunohistology, hemalaun counterstain. **C.** Staining with the anti-pan-RAV antibody shows the presence of abundant individual (arrowheads) and multiple IBs (short arrows) within hepatocytes. Again, a few larger IBs appear negative (large arrow). Immunohistology, hemalaun counterstain.

### Identification of reptarenaviruses, hartmaniviruses, and a chuvirus

To identify the infecting viruses, we isolated RNA from liver samples and performed a metatranscriptomic analysis, an approach we have successfully applied in earlier studies (9, 18, 22–24). We used the Basic Local Alignment Search Tool (BLAST, at https://blast.ncbi.nlm.nih.gov/Blast.cgi) to identify the viral sequences. The sequencing confirmed all snakes to be reptarenavirus infected, and similarly to earlier observations (8, 9, 22), all snakes harbored several L segments; however, we identified only a single S segment in each snake (Table 2). In addition to reptarenaviruses, the analysis revealed the presence of one hartmanivirus S and L segment pair in three of the four snakes (animals #1-3) studied (Table 2). We recovered complete coding sequences (CDSs) for one L and two S segments, and nearly complete CDS (covering >95% of the segment) of an additional hartmanivirus L segment. Genome *de novo* assembly using the sequence data obtained from these three snakes also produced close to identical contigs, varying in length from 10549 to 10718 nt (Table 2), that showed highest matches in the BLAST analysis to chuvirus-like viruses. Table 2 contains the virus names, contig lengths, GenBank accession numbers, and average coverages for the viruses identified, and Figure 4 shows the contig coverages nucleotide-by-nucleotide.

**Figure 4.**
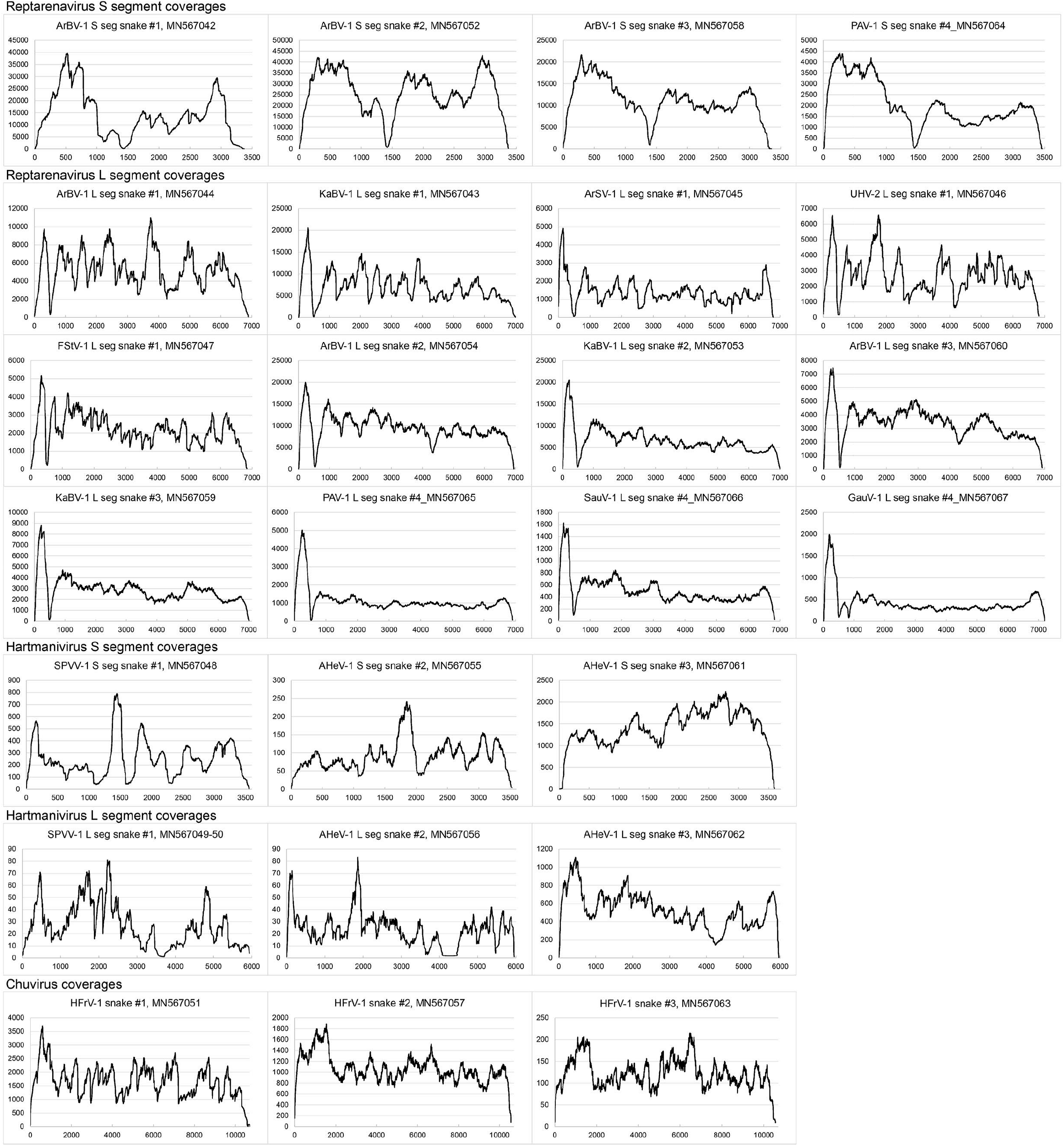
Coverages of the genomes and genome segments assembled. For all graphs: the y-axis represents the coverage and the x-axis indicates the nucleotide position.

Immunohistology served to detect reptarenavirus and hartmanivirus NP in cells with IB. Reptarenavirus NP expression was mainly restricted to the IBs (Figs. 2B and 3B, C), whereas hartmanivirus NP was also detected throughout the cytoplasm (Fig. 2C).

### Analysis of the identified reptarenavirus sequences

We used the PAirwise Sequence Comparison (PASC) web tool (available at https://www.ncbi.nlm.nih.gov/sutils/pasc/viridty.cgi?textpage=overview), recommended by the *Arenaviridae* study group of the International Committee on Taxonomy of Viruses (ICTV) for arenavirus classification (16), to analyze the identified reptarenavirus segments. The PASC results showed that we had recovered CDSs for seven novel L and two novel S segments (Table 2). The PASC analysis identified one of the L segments in snake #1 as UHV-2 (86.4% nucleotide, nt, identity), whereas in BLAST analysis three L segments in snakes #1-3 were >97% identical to Kaltenbach virus-1 (KaBV-1) (22), which is apparently not included in the PASC reference data set. Six L segments had less than 76% nucleotide identity to any currently known reptarenavirus sequences; Table 3 shows the nucleotide identity matrixes of the reptarenavirus segments. The analyses confirmed that we had recovered L segment CDSs for six novel reptarenavirus species (Tables 2 and 3): Aramboia boa virus-1 (ArBV-1, in animals #1-3), Arabuta Snake virus-1 (ArSV-1, in animal #1), Frankfurter Strasse virus-1 (FStV-1, in animal #1), Porto Alegre virus-1 (PAV-1, in animal #4), Saudades virus-1 (SauV-1, in animal #4), and Gaucho virus-1 (GauV-1, in animal #4). We found only a single S segment CDS for each of the studied snakes, and chose to name the S segments according to the L segment with highest coverage found in the same snake (Table 2): ArBV-1 (in animals #1-3) and PAV-1 (in animal #4). Phylogenetic analysis of the reptarenavirus L (Fig. 5) and S segments (Fig. 6A and B) showed the viruses to be distant from those present in GenBank, supporting their assignment as novel species. Notably, the reptarenavirus sequences recovered from the Brazilian snakes did not show geographical clustering but interspersed among the virus species detected from captive snakes in Europe and USA. As discussed previously (8), the phylogenetic trees constructed based on NP and GPC coding regions of reptarenavirus had incongruent topologies, suggesting recombination events to have occurred during the evolution of reptarenavirus S segments.

**Figure 5.**
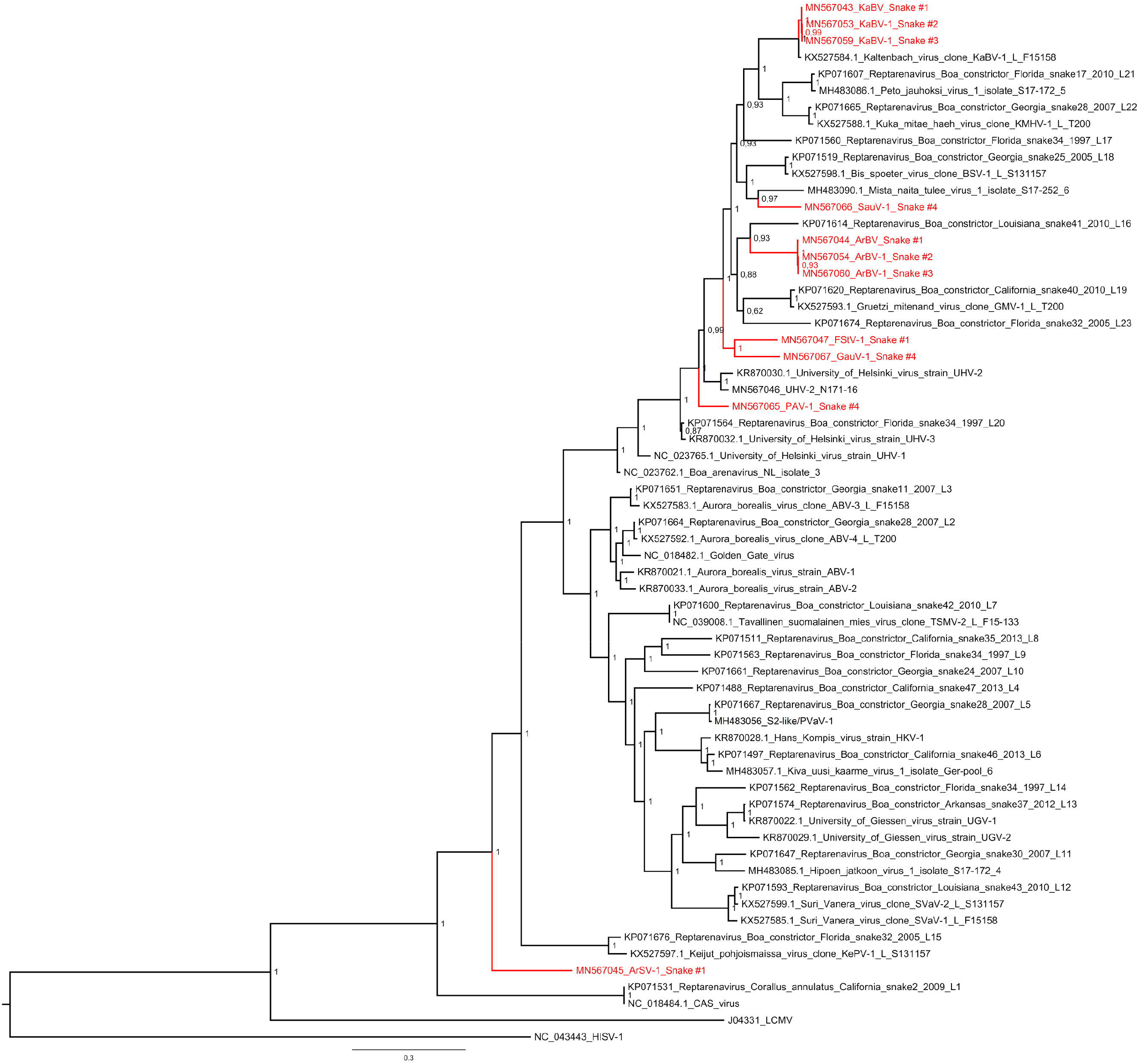
Maximum clade credibility tree of reptarenavirus L segments. The tree was constructed from amino acid sequences of the representatives of all reptarenavirus species and the strains identified in this study, using Bayesian MCMC method with Jones model of amino acid substitution. Posterior probabilities are shown in each node.

**Figure 6.**
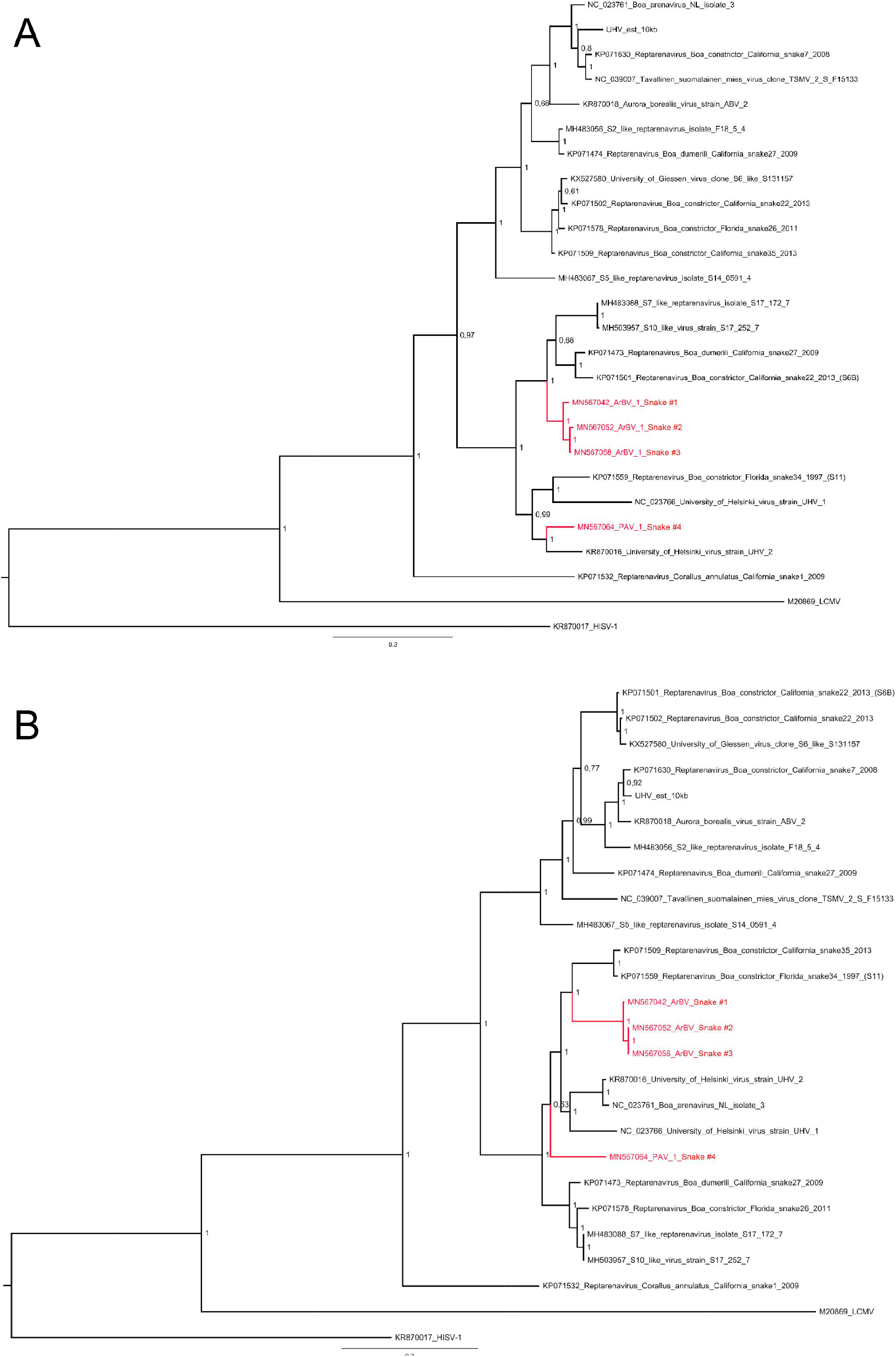
Maximum clade credibility trees of reptarenavirus GPCs and NPs. **A)** The phylogenetic tree based on the GPC amino acid sequences of the viruses identified in this study and those available in GenBank was constructed using Bayesian MCMC method with Blosum model of amino acid substitution. **B)** The phylogenetic tree based on the NP amino acid sequences of the viruses identified in this study and those available in GenBank was constructed using Bayesian MCMC method with Jones model of amino acid substitution.

### Analysis of the identified hartmanivirus sequences

We used the PASC tool also for analyzing the identified hartmanivirus S and L segment CDS, however, the analyses returned matches with very low sequence identities (22% and below, Table 2). To compare the sequences to known hartmaniviruses, we aligned the identified sequences with those found in the GenBank and generated nucleotide identity matrixes (Table 4). The analysis showed that the sequences are distant enough from each other and the known hartmaniviruses to represent new species: SetVetPat virus-1 (SPVV-1, in animal #1) and Andere Heimat virus-1 (AHeV-1, in animals #2 and #3). The phylogenetic analysis of hartmanivirus L and S segments suggested that these two viruses form a sister clade to the previously known hartmaniviruses (Fig. 7A-C).

**Figure 7.**
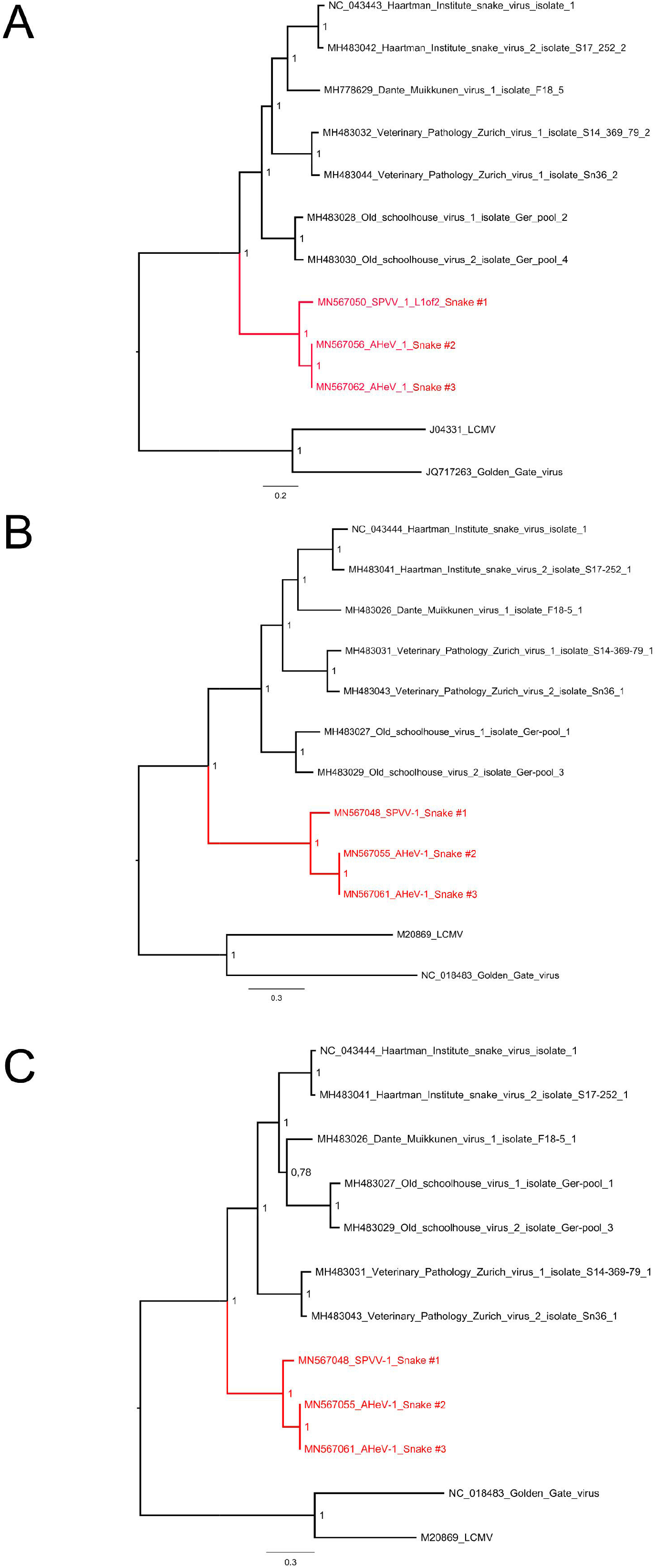
Maximum clade credibility trees for hartmanivirus RdRp, GPC, and NP. **A)** The phylogenetic tree based on the RdRp amino acid sequences of the viruses identified in this study and those available in GenBank was constructed using Bayesian MCMC method with Blosum model of amino acid substitution. **B)** The phylogenetic tree based on the GPC amino acid sequences of the viruses identified in this study and those available in GenBank was constructed using Bayesian MCMC method with Blosum model of amino acid substitution. **C)** The phylogenetic tree based on the NP amino acid sequences of the viruses identified in this study and those available in GenBank was constructed using Bayesian MCMC method with Wag model of amino acid substitution.

### Analysis of the novel species in the family *Chuviridae*

BLAST analysis identified three contigs that showed similarities to chuvirus-like viruses (family *Chuviridae*, genus *Mivirus*). These sequences had three ORFs in antigenomic orientation, representing the L, G, and N gene with RNA-dependent RNA polymerase (RdRp), glycoprotein (GP), and nucleoprotein (NP) as the respective protein products (Fig. 8A). We named the novel virus as Herr Frank virus-1 (HFrV-1, GenBank accession numbers in Table 2). BLAST analysis identified Guangdong red-banded snake chuvirus-like virus L protein (GenBank accession no. AVM87272.1) as the closest match (48.56% amino acid identity) for the HFrV-1 L gene (Fig. 8B) and putative GP (AVM87273.1) and NP (AVM87274.1) of as the closest matches for HFrV-1 GP and N gene (respective amino acid identities of 43.06% and 41.97%), Fig. 8B. In addition to BLAST analysis, we employed HMMSCAN (available at https://www.ebi.ac.uk/Tools/hmmer/search/hmmscan) to study the ORFs of HFrV-1. The analyses, presented in Fig. 8B, further confirmed the annotation of ORFs. The phylogenetic analysis of HFrV and the representatives of other chu-like virus RdRp sequences suggested that HFrV clusters together with Guangdong red-banded snake chuvirus-like virus and Wenling fish chu-like virus (25). Sanxia atyid shrip virus 4 formed an outgroup for these three vertebrate-associated chu-like viruses.

**Figure 8.**
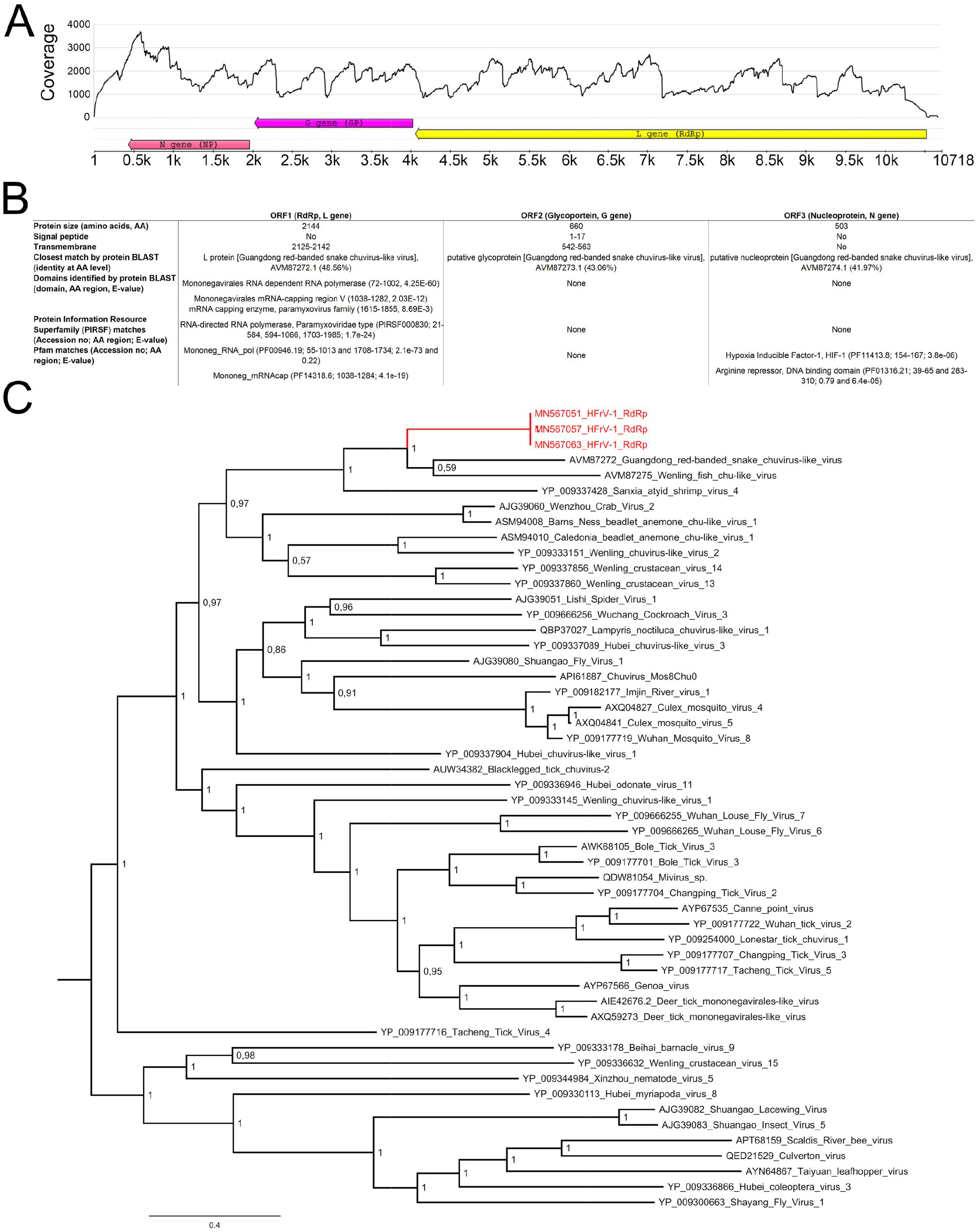
Genome organization, similarity analyses, and phylogenetic tree of HFrV-1. **A)** Genome organization and coverage (sequence from snake #1) of HFrV-1. The arrows represent the orientation of the open reading frames (ORFs). The L gene encodes RNA-dependent RNA polymerase (RdRp), G gene encodes glycoprotein (GP), and the N gene encodes nucleoprotein (NP). The coverage (y-axis) show sequencing depth at each nucleotide position (x-axis). **B)** Primary sequence, BLAST (https://blast.ncbi.nlm.nih.gov/Blast.cgi) and HMMSCAN (https://www.ebi.ac.uk/Tools/hmmer/) analysis of the HFrV-1 ORFs. **C)** A maximum clade credibility tree based on the RdRp amino acid sequences of chu-like and chuviruses. The tree was constructed using Bayesian MCMC method with Blosum model of amino acid substitution. Posterior probabilities are shown in each node.

## DISCUSSION

This study aimed to confirm the presence of BIBD and reptarenaviruses in *Boa constrictors*, indigenous to the Brazilian wildlife. We initially used immunohistology to confirm the presence of reptarenavirus-induced IBs in all four studied snakes, thus confirming the BIBD diagnosis. The subsequent metatranscriptomic analysis of the livers confirmed the presence of reptarenaviruses, and helped to identify two novel hartmanivirus species and a novel chuvirus in the snakes. The affected boas had lived in captivity, with contact to other snakes. However, three of the snakes originated from the Amazon region in Brazil, which could indicate that both reptarenaviruses and hartmaniviruses exist in wild snakes in this region.

At the time of sampling/euthanasia, all snakes suffered from bacterial (and fungal) infections, caused by opportunistic agents that are part of the mucosal flora in various species including snakes. These agents occur in the oral cavities of captive snakes with stomatitis or in snakes with septicemia (26, 27); some, like *Enterobacter gergoviae*, exist in water or soil (28). All can cause disease, particularly in immunocompromised patients. Secondary infections with related inflammatory processes are common in snakes with BIBD, indicating an immunosuppressive effect at least in later, chronic stages of BIBD. Much less is known about the effect of acute reptarenavirus infections. Based on our experience with *in vitro* studies, we assume that larger, often more amphophilic than eosinophilic IBs represent late stages of IB formation. Such IBs often exhibit only a peripheral positive immunohistological reaction for viral NP, similarly to the reaction seen in the tissues of the snakes in this study. *In vitro*, small irregular IBs appear at early stages (approx. 3 days post infection, dpi) of reptarenavirus infection, and from 12 dpi onwards the IBs become larger and more electron dense (unpublished data). Thus, we assume that the snakes included in this study were chronically reptarenavirus infected, and therefore immunocompromised (the hartmanivirus and a chuvirus co-infections may also have contributed), which in turn led to the observed secondary infections.

Metatranscriptomic analysis of the animals revealed the presence of multiple reptarenavirus L segments, but only a single S segment per snake. The finding is very surprising, since others and we have observed reptarenavirus co-infections to be common in snakes with BIBD (8, 9). One of the identified segments, the KaBV-1 L segment (animals #1-3), showed a striking 97% identity to a previously identified (22) reptarenavirus genome segment. In addition, one L segment showed approximately 86% identity to the UHV-2 L segment, but the other segments differed enough from the previously identified reptarenavirus segments to warrant classification as novel reptarenavirus species. Interestingly, we did not find University of Giessen virus or “S6-like” S segment in any of the studied snakes, even though the segment is most often reported in captive snakes with BIBD (8, 9, 13, 22). In the phylogenetic trees, most of the reptarenaviruses identified from the Brazilian snakes interspersed among the virus species previously detected from captive snakes in Europe and USA. Notably, one of the reptarenavirus species, KaBV-1, showed high identity to a strain identified from captive snakes in Switzerland (22). This may indicate that reptarenaviruses have been introduced to captive snake populations by wild boas and then been exported with them, due to the lack of clinical signs of infection/disease (13). A great number of snakes are traded annually, according to CITES 20,000 snakes (6,600 pythonids and 3,100 boids) were transported in 2018 alone (https://trade.cites.org/en/cites_trade/#). While the trading and transport may technically follow CITES regulations, a great number of transported animals are likely not captive bred but wild-caught. In addition to the transport of animals following CITES regulations, smuggling of wild-caught snakes occurs frequently. The spread would further be aided by the fact that reptarenavirus infection does not induce clinical signs rapidly, especially not in boas (10), and is vertically transmitted (22). The studies on snakes with BIBD strongly suggest that reptarenavirus L and S segments are able to pair with each other rather freely, since most often the individuals harbor more L than S segments (8, 9, 9). Assuming that snakes, or better boas and pythons, are the reservoir hosts of reptarenaviruses and that reptarenaviruses have co-evolved with their reservoir hosts, then multiple crossspecies transmission events could explain the status quo in captive snakes. However, with the current set of data we cannot rule out the possibility that the wild-caught boas included in the study had not been infected during co-housing.

The identification of novel hartmanivirus species in Brazilian *B. constrictor* snakes is interesting, since up to now hartmanivirus infection has only been reported in European captive snakes (9, 18). The hartmanivirus infected snakes included in the present study had developed BIBD as confirmed by the presence of IBs in both blood smear and tissues, which is in accordance with our earlier findings (9, 18). They showed strong expression of hartmanivirus NP in parenchymal cells in various organs. We have thus far detected hartmaniviruses mainly in snakes with BIBD, however, the fact that we mainly look for viruses in diagnostic cases might introduce a bias and could explain the seeming correlation between hartmanivirus infection and BIBD. In fact, when studying samples collected from a single breeding colony for the presence of IBs, reptarenaviruses, and hartmaniviruses, we did not find a significant correlation between hartmanivirus infection and BIBD (13). Although hartmanivirus infection appears to most often occur simultaneously to reptarenavirus infection, hartmaniviruses do can infect and replicate without a co-infecting reptarenavirus (18) and further studies need to address their pathogenicity. Alike reptarenaviruses, the origin of hartmaniviruses remains unknown. In addition to snakes being the reservoir hosts of the viruses, one could speculate that blood-feeding parasites e.g. mites, ticks, mosquitoes, etc. would serve as reservoirs and/or vectors in virus transmission.

The novel chuvirus, HFrV-1, found in three BIBD positive snakes originating from a snake sanctuary in the Amazonas region, but housed in a smaller colony for several years afterwards, was an unexpected finding. By amino acid identity, the closest relative to the newly found mononegavirus is the Guandong red-banded snake Chuvirus, which was identified from a liver sample of a Chinese snake (25). In general, the identification of chu-like viruses in fish and snakes from different continents (Asia and the Americas) suggests that chuviruses might be common and geographically widespread. Due to bacterial and viral co-infections in the snakes with HFrV-1 infection, we cannot draw conclusions on the potential morbidity of HFrV-1. The fact that the identified viruses showed nearly identical sequences suggests that the Chuvirus infection may have occurred during captivity.

According to Whitfield et al, the major threats of declining reptilian populations are habitat loss and degradation, introduction of invasive species, environmental pollution, disease, unsustainable use and global climate change (29). Our results raise an obvious question: From where did the reptarenaviruses come that infected the diseased snakes? Both co-housed imported snakes and local wild snakes are a potential source of infection. Animals #1-3 originated from the Amazonas region, where they resided in a snake sanctuary before moving to a private collection in Porto Alegre. It was not possible to obtain information on other snake species housed in the sanctuary, since it was closed several years back. We also lack more specific information on the origin of the snakes. Similarly, the origin of animal #4 remained unknown. However, all animals studied were *B. constrictor*, indigenous to Brazil, and it is therefore possible that they originated from the wild. If snakes are not the reservoir hosts of reptarenaviruses, then the occurrence of BIBD-positive *B. constrictor* in Brazil is an alarming signal posing a potential threat to Brazilian wild *B. constrictor* populations.

## MATERIALS AND METHODS

### Animals

The study was undertaken on four captive adult *B. constrictor constrictor* snakes. Three derived from a single private owner (animals #1-3), the fourth from a zoological garden (Table 1). All animals were submitted for diagnostic post mortem examination to the Department of Veterinary Pathology in Porto Alegre. Tissue specimens were fixed in 10% buffered formalin for histological and immunohistological examination. Additional sets of samples were stored frozen in RNAlater™ Stabilization Solution (ThermoFischer Scientific) for RNA extraction. For animals #1, #3 and #4, samples from the oral and nasal lesions, respectively, were subjected to a routine bacteriological examination; for animal #1, a routine mycological examination was also performed.

### Histology and immunohistology

Formalin-fixed tissue specimens were trimmed and routinely paraffin wax embedded. Consecutive sections (3-5 μm) were prepared and stained with hematoxylin-eosin (HE) and special stains (Periodic acid-Schiff (PAS) reaction, Grocott methenamine silver stain), when appropriate. Further sections were subjected to immunohistological staining for reptarenavirus and hartmanivirus NP as described (5).

### Antibodies, protein expression, and immunization

The anti-UHV NP and anti-UHV NP C-terminus antibodies were described earlier (5, 30). To generate broadly cross-reactive antiserum against reptarenavirus NPs, we performed amino acid alignment for the reptarenavirus NP available in GenBank. Based on homology between the sequences, we selected the following regions: amino acids 47-140 from UGV-1 (GenBank: YP_009508464.1), 173-224 from UHV-1 (YP_009019205.1), 233-270 from UHV-1 (YP_009019205.1), 286-359 from UGV-1 (YP_009508464.1), 208-280 from UGV-1 (YP_009508464.1), and 494-567 from UHV-1 (YP_009019205.1). To generate a cross-reactive antiserum against hartmanivirus NPs, we used the same approach and selected the following regions: amino acids 199-256 from Veterinary Pathology Zürich virus-1, VPZV-1 (AZI72586.1); 132-180 from Haartman Institute Snake virus-2, HISV-2 (AZI72594.1); 257-299 from VPZV-1 (AZI72586.1), and 312-364 from VPZV-2 (AZI72596.1). We included five glycine residues between the selected epitopes, and ordered the engineered proteins as synthetic genes optimized for *E. coli* expression in pET-20b(+) plasmid from GenScript. We transformed One Shot™ BL21(DE3) Chemically Competent (Thermo Scientific) *E. coli* with the plasmids following the manufacturer’s protocol, and performed protein expression and purification via his-tag as described (23, 30). Antisera against the purified proteins were raised by BioGenes, as described in earlier studies (23, 24, 30). We designated the novel antisera as anti-pan-RAV and anti-pan-hartmani.

### Next generation sequencing (NGS) and genome assembly

We extracted RNAs for NGS from liver samples stored frozen in RNAlater™ (the sample from animal #1 had been kept at ambient temperature for a few weeks prior to extraction) as described (22), prepared NGS libraries, and performed sequencing and subsequent genome assembly as described (22, 23).

### Reverse transcriptase-polymerase chain reaction (RT-PCR) and Sanger sequencing

For the sample that had been stored in RNAlater at ambient temperature (animal #1) we obtained open reading frames (ORFs) for several NP, ZP, GPC, and RdRp genes instead of complete L and S segments by NGS and de novo assembly. To complete the L and S segment sequences, we designed the following primers to amplify the missing intergenic regions: Br_GPC1 5’-ACACTTGGATTCTATGGGAGT-3’, Br_NP1 5’-ACTGCATGGTGTTCTCAAG-3’, Br_ZP1 5’-GAGTCTAACCAATCCCAGAA-3’, Br_ZP2 5’-CATGCCTAATGGCAAAAC-3’, Br_ZP3 5’-CAGAATGTAGGGCAACAC-3’, Br_ZP4 5’-AGGGTCTAAATCAACATCCC-3’, Br_UHV_RdRp 5’-GTCAGAATATCACTCCTGGAG-3’, Br_RdRp2 5’-TAGGGTGACACTTTTGAAGG-3’, Br_RdRp3 5’-GAACATTAGGGTATCACTCCTC-3’, and Br_RdRp4 5’-AGAGTCTAAGGGTCCTGGA-3’. We performed RT-PCR with all primer combinations (ZPs with RdRps, and GPC with NP) as described in (22), and used GeneJET Gel Extraction Kit (ThermoFisher Scientific) to purify the RT-PCR products which were further cleaned by AMPure XP beads (Beckman Coulter) before ligation to plasmid using the Zero Blunt TOPO PCR Cloning Kit (ThermoFisher Scientific); all steps were according to the manufacturer’s instructions. Chemically competent *E. coli* (TOP10, ThermoFisher Scientific) transformed with the ligated plasmids were grown on LB plates with 100 μg/ml of ampicillin O/N at 37 °C, colonies were picked and grown in 5 ml of 2x YT medium with 100 μg/ml of ampicillin O/N at 37 °C. The plasmids were purified from 2 ml of the O/N culture using the GeneJET Plasmid Miniprep Kit (ThermoFisher Scientific), and sent for Sanger sequencing to DNA Sequencing and Genomics, Institute of Biotechnology, University of Helsinki.

### Phylogenetic analysis

The amino acid sequences of the representatives of all reptarena- and hartmanivirus species were downloaded from the GenBank, and aligned with the amino acid sequences of viruses identified in this study using the MAFFT E-INS-i algorithm (31). For chuvirus-like sequences, 100 closest BLASTx matches for the putative RdRp amino acid sequence were downloaded from GenBank and aligned as indicated above. Redundant sequences (fragmental and identical sequences) were removed from the dataset.

The best-fit amino acid substitution models and phylogenetic trees were inferred using the Bayesian Monte Carlo Markov Chain (MCMC) method implemented in MrBayes v3.2.6. (32). MrBayes was run for 500,000 generations and sampled every 5,000 steps, with final standard deviations between two runs < 0.02 for all analyses. The analyses were carried out at the CSC server (IT Center for Science Ltd., Espoo, Finland).

### Data availability

The names for newly sequenced viruses with corresponding abbreviations and GenBank accession numbers are provided in Table 2.

## ACKNOWLEDGEMENTS

The authors are grateful to the laboratory staff of the Histology Laboratory, Institute of Veterinary Pathology, Vetsuisse Faculty, University of Zürich, and colleagues and professors of Universidade Federal do Rio Grande do Sul for excellent technical support. They also wish to thank L Sonne and SP Pavarini who assisted in the snake necropsies during their undergraduate studies. The study received financial support from the Leading House for the Latin American Region, University of St. Gallen, Switzerland, and the Academy of Finland (grant numbers 1308613 and 1314119). This work is dedicated to the memory of Dr. Herbert Frank and to his colleagues at the Institute of Veterinary Pathology, Faculty of Veterinary Medicine, Justus-Liebig-University Giessen, Germany, where the three senior authors of this publication were trained in veterinary pathology.

---

| **Snake No.** | **Virus name (abbreviation)** | **Segment** | **Contig length** | **GenBank accession no.** | **Average coverage** | **Closest match by PASC (% identity, GenBank accession no.)** |
| --- | --- | --- | --- | --- | --- | --- |
| **1** | Aramboia boa virus-1 (ArBV-1) | S | 3372 | MN567042 | 14173.9 | 74.0%. KP071473.1 Unclassified Reptarenavirus |
| **1** | Aramboia boa virus-1 (ArBV-1) | L | 6883 | MN567044 | 5232.6 | 72.7%. KR870030.1 University of Helsinki reptarenavirus |
| **1** | Kaltenbach virus-1 (KaBV-1) | L | 7010 | MN567043 | 7105.8 | 72.4%. KP071677.1 Unclassified Reptarenavirus |
| **1** | Arabuta Snake virus-1 (ArSV-1) | L | 6780 | MN567045 | 1413.6 | 63.3%. KR870020.1 University of Helsinki reptarenavirus |
| **1** | University of Helsinki virus-2 (UHV-2) | L | 6813 | MN567046 | 2765.8 | 86.4%. KR870030.1 University of Helsinki reptarenavirus |
| **1** | Frankfurter Strasse virus-1 (FStV-1) | L | 6831 | MN567047 | 2233.2 | 74.1%. KR870030.1 University of Helsinki reptarenavirus |
| **1** | SetPatVet virus-1 (SPVV-1) | S | 3560 | MN567048 | 268.3 | 21.8%. FJ607031.1 Lymphocytic choriomeningitis mammarenavirus |
| **1** | SetPatVet virus-1 (SPVV-1) | L | 3578 and 2223 | MN567049 and MN567050 | 35.1 | N.A. |
| **1** | Herr Frank virus-1 (HFrV-1) | N.A. | 10718 | MN567051 | 1698.9 | N.A. |
| **2** | Aramboia boa virus-1 (ArBV-1) | S | 3372 | MN567052 | 26110.1 | 73.7%. KP071473.1 Unclassified Reptarenavirus |
| **2** | Aramboia boa virus-1 (ArBV-1) | L | 6948 | MN567054 | 9586.7 | 72.8%. KR870030.1 University of Helsinki reptarenavirus |
| **2** | Kaltenbach virus-1 (KaBV-1) | L | 7009 | MN567053 | 6688.2 | 72.3%. KP071567.1 Unclassified Reptarenavirus |
| **2** | Andere Heimat virus-1 (AHeV-1) | S | 3534 | MN567055 | 99.3 | 22.0%. FJ607031.1 Lymphocytic choriomeningitis mammarenavirus |
| **2** | Andere Heimat virus-1 (AHeV-1) | L | 5953 | MN567056 | 28.5 | 18.7%. MG812678.1 Lassa mammarenavirus |
| **2** | Herr Frank virus-1 (HFrV-1) | N.A. | 10549 | MN567057 | 1073.9 | N.A. |
| **3** | Aramboia boa virus-1 (ArBV-1) | S | 3351 | MN567058 | 11236.5 | 74.0%. KP071671.1 Unclassified Reptarenavirus |
| **3** | Aramboia boa virus-1<br>(ArBV-1) | L | 6926 | MN567060 | 3443.9 | 72.7%. KR870030.1 University of Helsinki reptarenavirus |
| **3** | Kaltenbach virus-1<br>(KaBV-1) | L | 7007 | MN567059 | 2913.8 | 72.2%. KR870030.1 University of Helsinki reptarenavirus |
| **3** | Andere Heimat virus-1<br>(AHeV-1) | S | 3598 | MN567061 | 1433.3 | 21.8%. NC_004294.1 Lymphocytic choriomeningitis mammarenavirus |
| **3** | Andere Heimat virus-1<br>(AHeV-1) | L | 5961 | MN567062 | 548.3 | 18.9%. MG812678.1 Lassa mammarenavirus |
| **3** | Herr Frank virus-1<br>(HFrV-1) | N.A. | 10574 | MN567063 | 126.8 | N.A. |
| **4** | Porto Alegre virus-1<br>(PAV-1) | S | 3464 | MN567064 | 2071.6 | 73.3%. KP071559.1 Unclassified Reptarenavirus |
| **4** | Porto Alegre virus-1<br>(PAV-1) | L | 6911 | MN567065 | 1158.3 | 79.6%. KP071549.1 Unclassified Reptarenavirus |
| **4** | Saudades virus-1<br>(SauV-1) | L | 6835 | MN567066 | 523.1 | 73.2%. KP071479.1 Unclassified Reptarenavirus |
| **4** | Gaucho virus-1<br>(GauV-1) | L | 7203 | MN567067 | 438.3 | 72.8%. KR870030.1 University of Helsinki reptarenavirus |

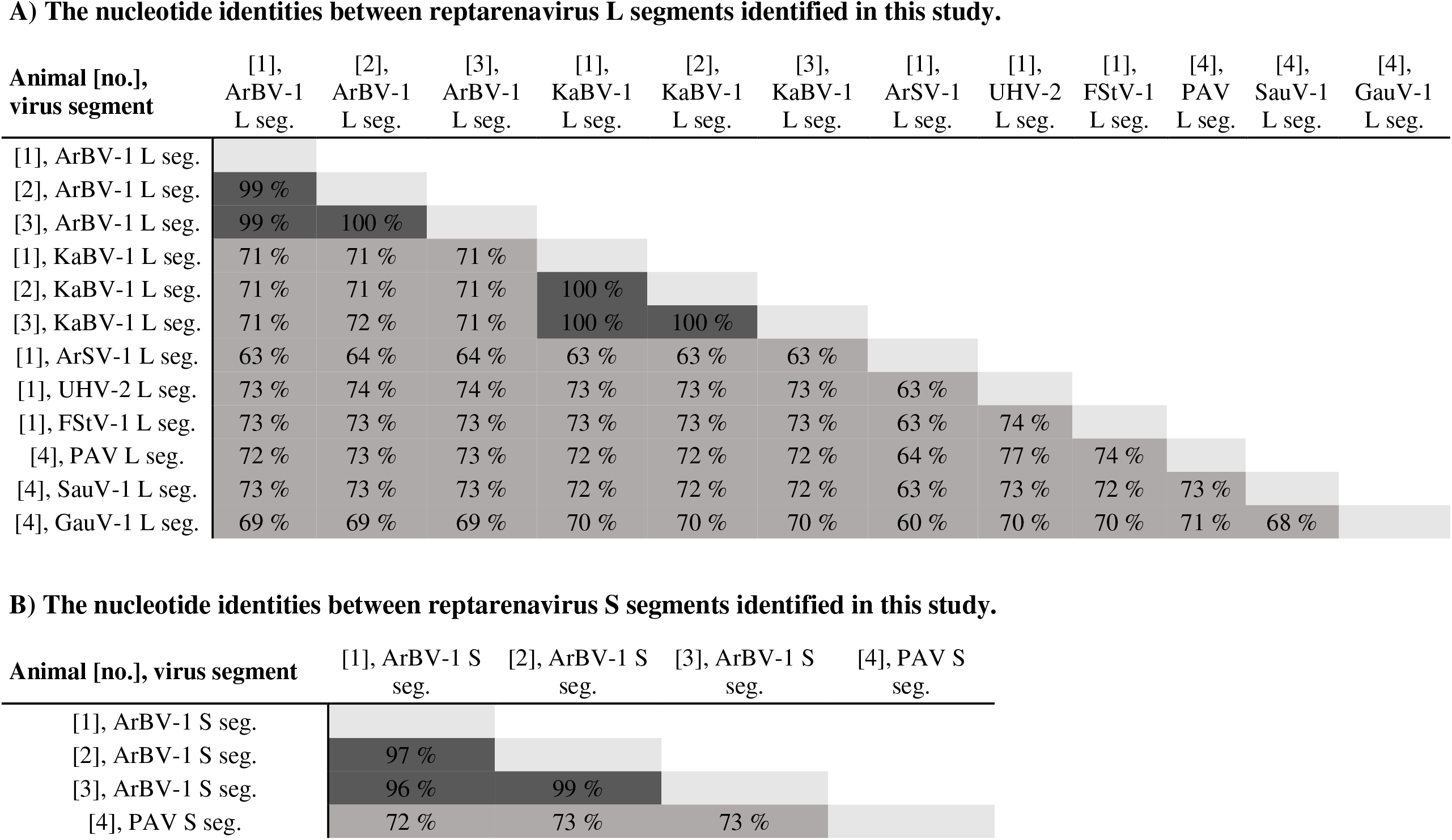

**A) The nucleotide identities between reptarenavirus L segments identified in this study.**
| **Animal [no.],**<br>**virus segment** | [1],<br>ArBV-1<br>L seg. | [2],<br>ArBV-1<br>L seg. | [3],<br>ArBV-1<br>L seg. | [1],<br>KaBV-1<br>L seg. | [2],<br>KaBV-1<br>L seg. | [3],<br>KaBV-1<br>L seg. | [1],<br>ArSV-1<br>L seg. | [1],<br>UHV-2<br>L seg. | [1],<br>FStV-1<br>L seg. | [4],<br>PAV<br>L seg. | [4],<br>SauV-1<br>L seg. | [4],<br>GauV-1<br>L seg. |
| --- | --- | --- | --- | --- | --- | --- | --- | --- | --- | --- | --- | --- |
| [1], ArBV-1 L seg. |  |  |  |  |  |  |  |  |  |  |  |  |
| [2], ArBV-1 L seg. | 99 % |  |  |  |  |  |  |  |  |  |  |  |
| [3], ArBV-1 L seg. | 99 % | 100 % |  |  |  |  |  |  |  |  |  |  |
| [1], KaBV-1 L seg. | 71 % | 71 % | 71 % |  |  |  |  |  |  |  |  |  |
| [2], KaBV-1 L seg. | 71 % | 71 % | 71 % | 100 % |  |  |  |  |  |  |  |  |
| [3], KaBV-1 L seg. | 71 % | 72 % | 71 % | 100 % | 100 % |  |  |  |  |  |  |  |
| [1], ArSV-1 L seg. | 63 % | 64 % | 64 % | 63 % | 63 % | 63 % |  |  |  |  |  |  |
| [1], UHV-2 L seg. | 73 % | 74 % | 74 % | 73 % | 73 % | 73 % | 63 % |  |  |  |  |  |
| [1], FStV-1 L seg. | 73 % | 73 % | 73 % | 73 % | 73 % | 73 % | 63 % | 74 % |  |  |  |  |
| [4], PAV L seg. | 72 % | 73 % | 73 % | 72 % | 72 % | 72 % | 64 % | 77 % | 74 % |  |  |  |
| [4], SauV-1 L seg. | 73 % | 73 % | 73 % | 72 % | 72 % | 72 % | 63 % | 73 % | 72 % | 73 % |  |  |
| [4], GauV-1 L seg. | 69 % | 69 % | 69 % | 70 % | 70 % | 70 % | 60 % | 70 % | 70 % | 71 % | 68 % |  |

**B) The nucleotide identities between reptarenavirus S segments identified in this study.**
| **Animal [no.], virus segment** | [1], ArBV-1 S<br>seg. | [2], ArBV-1 S<br>seg. | [3], ArBV-1 S<br>seg. | [4], PAV S<br>seg. |
| --- | --- | --- | --- | --- |
| [1], ArBV-1 S seg. |  |  |  |  |
| [2], ArBV-1 S seg. | 97 % |  |  |  |
| [3], ArBV-1 S seg. | 96 % | 99 % |  |  |
| [4], PAV S seg. | 72 % | 73 % | 73 % |  |

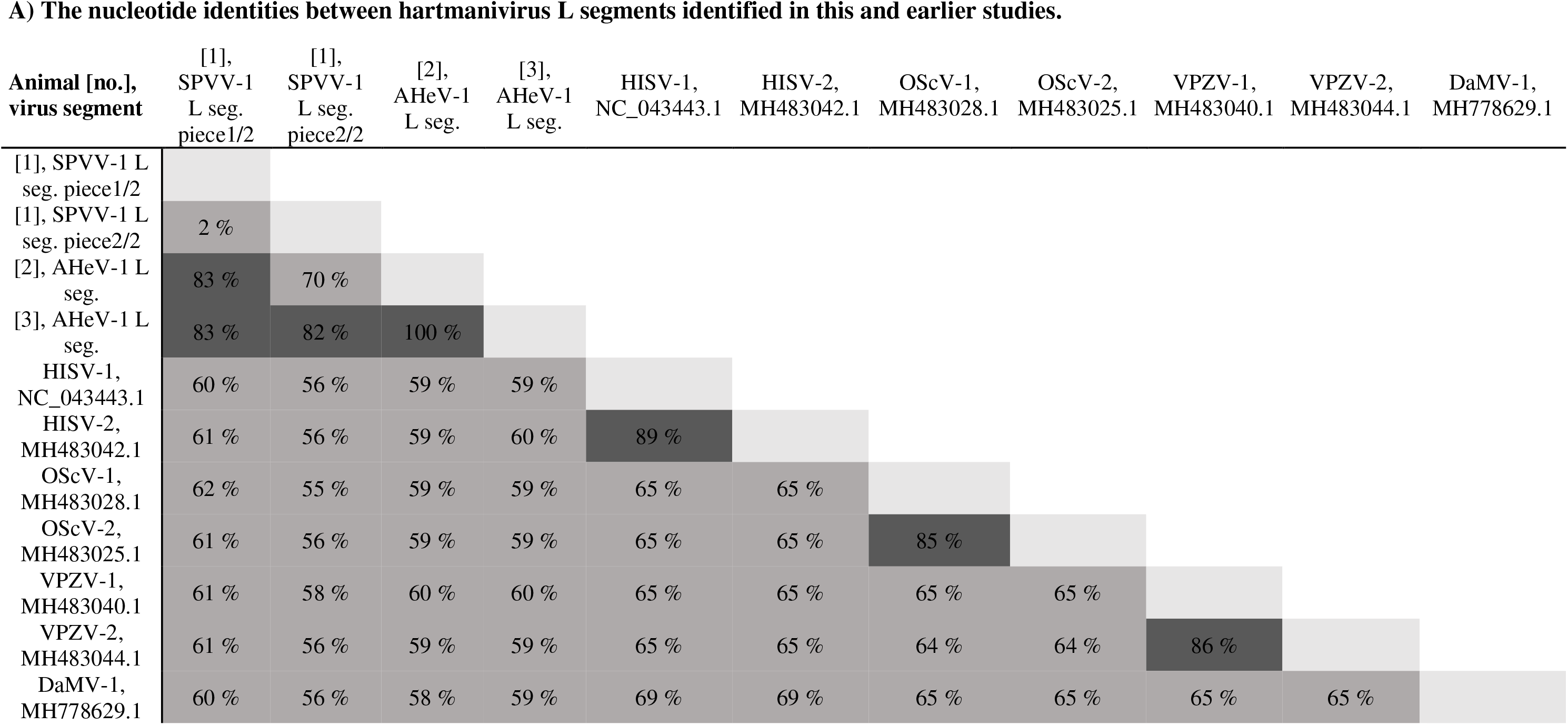
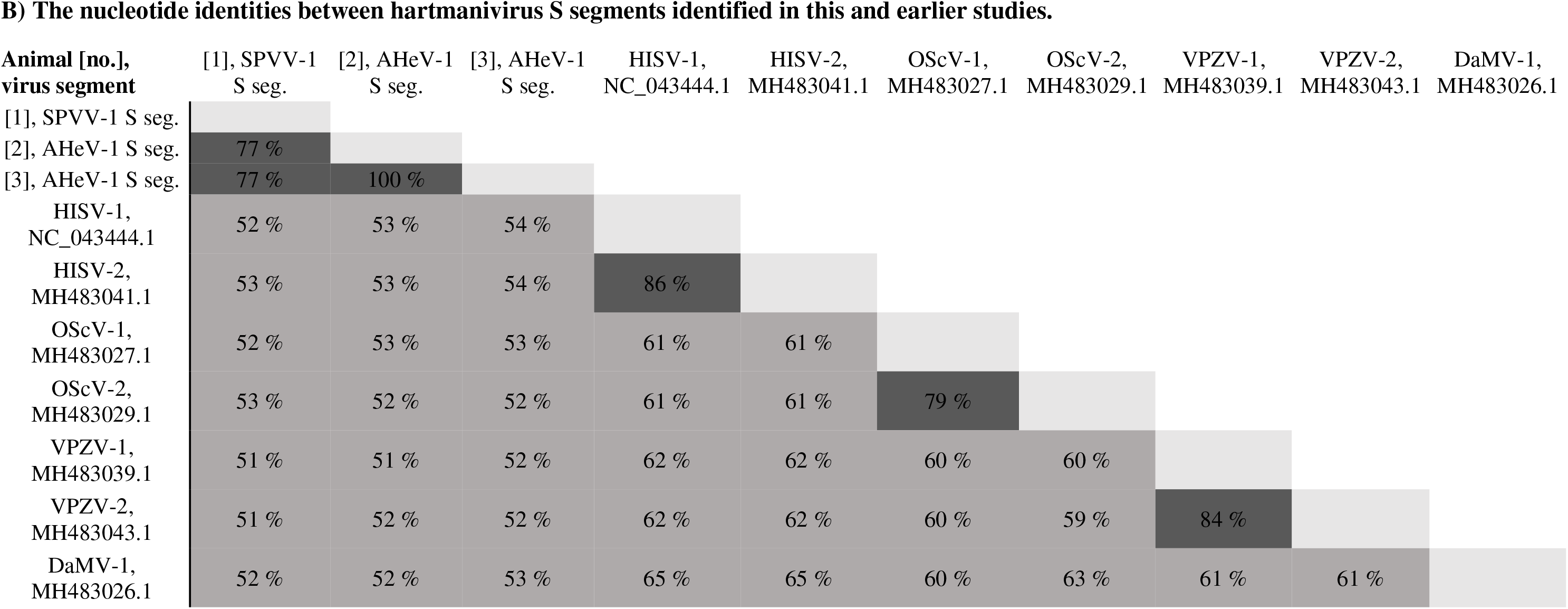

**A) The nucleotide identities between hartmanivirus L segments identified in this and earlier studies.**
| **Animal [no.],**<br>**virus segment** | [1],<br>SPVV-1<br>L seg.<br>piece1/2 | [1],<br>SPVV-1<br>L seg.<br>piece2/2 | [2],<br>AHeV-1<br>L seg. | [3],<br>AHeV-1<br>L seg. | HISV-1,<br>NC_043443.1 | HISV-2,<br>MH483042.1 | OScV-1,<br>MH483028.1 | OScV-2,<br>MH483025.1 | VPZV-1,<br>MH483040.1 | VPZV-2,<br>MH483044.1 | DaMV-1,<br>MH778629.1 |
| --- | --- | --- | --- | --- | --- | --- | --- | --- | --- | --- | --- |
| [1], SPVV-1 L<br>seg. piece1/2 |  |  |  |  |  |  |  |  |  |  |  |
| [1], SPVV-1 L<br>seg. piece2/2 | 2 % |  |  |  |  |  |  |  |  |  |  |
| [2], AHeV-1 L<br>seg. | 83 % | 70 % |  |  |  |  |  |  |  |  |  |
| [3], AHeV-1 L<br>seg. | 83 % | 82 % | 100 % |  |  |  |  |  |  |  |  |
| HISV-1,<br>NC_043443.1 | 60 % | 56 % | 59 % | 59 % |  |  |  |  |  |  |  |
| HISV-2,<br>MH483042.1 | 61 % | 56 % | 59 % | 60 % | 89 % |  |  |  |  |  |  |
| OScV-1,<br>MH483028.1 | 62 % | 55 % | 59 % | 59 % | 65 % | 65 % |  |  |  |  |  |
| OScV-2,<br>MH483025.1 | 61 % | 56 % | 59 % | 59 % | 65 % | 65 % | 85 % |  |  |  |  |
| VPZV-1,<br>MH483040.1 | 61 % | 58 % | 60 % | 60 % | 65 % | 65 % | 65 % | 65 % |  |  |  |
| VPZV-2,<br>MH483044.1 | 61 % | 56 % | 59 % | 59 % | 65 % | 65 % | 64 % | 64 % | 86 % |  |  |
| DaMV-1,<br>MH778629.1 | 60 % | 56 % | 58 % | 59 % | 69 % | 69 % | 65 % | 65 % | 65 % | 65 % |  |

**B) The nucleotide identities between hartmanivirus S segments identified in this and earlier studies.**
| **Animal [no.],**<br>**virus segment** | **[1], SPVV-1**<br>**S seg.** | **[2], AHeV-1**<br>**S seg.** | **[3], AHeV-1**<br>**S seg.** | **HISV-1,**<br>**NC_043444.1** | **HISV-2,**<br>**MH483041.1** | **OScV-1,**<br>**MH483027.1** | **OScV-2,**<br>**MH483029.1** | **VPZV-1,**<br>**MH483039.1** | **VPZV-2,**<br>**MH483043.1** | **DaMV-1,**<br>**MH483026.1** |
| --- | --- | --- | --- | --- | --- | --- | --- | --- | --- | --- |
| [1], SPVV-1 S seg. |  |  |  |  |  |  |  |  |  |  |
| [2], AHeV-1 S seg. | 77 % |  |  |  |  |  |  |  |  |  |
| [3], AHeV-1 S seg. | 77 % | 100 % |  |  |  |  |  |  |  |  |
| HISV-1,<br>NC_043444.1 | 52 % | 53 % | 54 % |  |  |  |  |  |  |  |
| HISV-2,<br>MH483041.1 | 53 % | 53 % | 54 % | 86 % |  |  |  |  |  |  |
| OScV-1,<br>MH483027.1 | 52 % | 53 % | 53 % | 61 % | 61 % |  |  |  |  |  |
| OScV-2,<br>MH483029.1 | 53 % | 52 % | 52 % | 61 % | 61 % | 79 % |  |  |  |  |
| VPZV-1,<br>MH483039.1 | 51 % | 51 % | 52 % | 62 % | 62 % | 60 % | 60 % |  |  |  |
| VPZV-2,<br>MH483043.1 | 51 % | 52 % | 52 % | 62 % | 62 % | 60 % | 59 % | 84 % |  |  |
| DaMV-1,<br>MH483026.1 | 52 % | 52 % | 53 % | 65 % | 65 % | 60 % | 63 % | 61 % | 61 % |  |

